# A novel SXXLF motif in the FXR N-terminal domain mediates coregulator and interdomain interactions

**DOI:** 10.64898/2026.05.18.724725

**Authors:** Priscilla Villalona, Thilini Pulahinge, Tracy Yu, Jordan Wenning, Crawford Joseph Frisbie, Jill Magafas, C. Denise Okafor

## Abstract

The nuclear receptor superfamily is comprised of ligand-regulated transcription factors that contain an intrinsically disordered domain at the amino-terminal end, known as the N-terminal domain (NTD). While this poorly conserved domain is known to possess ligand-independent activation function (AF-1), few NTD functions are conserved between nuclear receptors (NRs). Identified roles in other receptors include androgen receptor (AR), estrogen receptor (ER) and mineralocorticoid receptor (MR). Here, we aim to define the function of the NTD of the farnesoid X receptor (FXR), a crucial regulator of lipid and bile acid metabolism. We show that the NTD engages in interdomain contact with other FXR domains. We also observe that the NTD interacts directly with coregulator proteins. Using mutagenesis, mammalian two-hybrid assays and molecular dynamics simulations, we identify and validate a novel SXXLF motif in the NTD which mediates interactions with both coregulators and the ligand binding domain. Mutation of the motif induces large changes in conformational and allosteric coupling in FXR. Our study identifies a new nuclear receptor-interacting motif that modulates the transcriptional activity of FXR.

**Graphical Abstract:** FXR-NTD regulates transcriptional activity through interdomain communication with the LBD and is also involved in co-activator recruitment. The SENLF motif is the first defined functional element within the FXR-NTD and mediates both NTD-LBD interaction and selective co-activator engagements to drive NTD-mediated transcriptional activity.

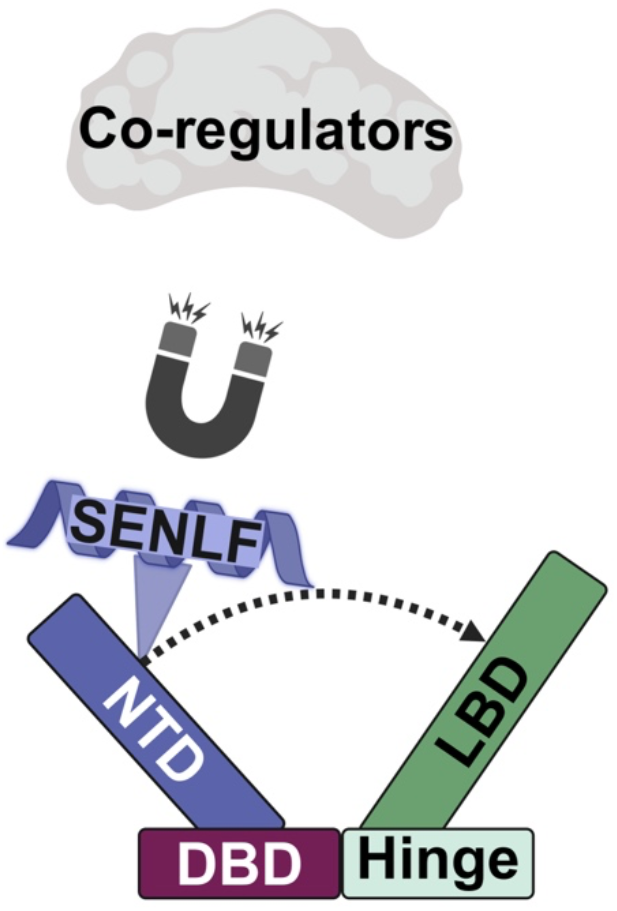

## Introduction

The FXR (NR1H4) is a member of the nuclear receptor superfamily and is regulated by bile acids.^[1]^ FXR is involved in a wide array of biological processes, including bile acid homeostasis, glucose metabolism, and lipid metabolism.^[2–4]^ Like other NRs, FXR has a modular structure composed of various domains: the NTD, the DNA-binding domain (DBD), a flexible hinge region, and lastly the ligand-binding domain (LBD).^[5,6]^ The NTD is a predominantly disordered domain in NRs, including FXR, and includes an AF-1 region where coregulators can bind.^[5,6]^ Following the NTD is the DBD, which contains two zinc-coordinated fingers that interact with specific DNA sequences known as FXR response elements (FXRE) recognized by the DBD.^[5,6]^ The LBD is usually formed by 11 helices and 4 beta strands that arrange to create a three-layer α-helical sandwich structure, with the hydrophobic amino acids at the core comprising the ligand-binding pocket (LBP).^[5,6]^ The last helix (H12) also contains the activation function 2 (AF-2) surface, which plays a role in ligand-induced coregulator recruitment, along with H3 and H4.^[5,6]^ For FXR, many studies have characterized the LBD and DBD of the receptor, expanding our understanding of their functions.^[7,8]^

In contrast, the NTD in FXR and other NRs is highly disordered, making it challenging to use in structural studies to understand its function. In other NRs, their NTDs have been found to participate in binding with other binding partners and engage in interdomain crosstalk.^[9]^ In the estrogen receptor, the AF-1 region, which is a site for ligand-independent coregulator binding, has been found to interact with GRIP1 and SRC1α, which are members of the p160 coactivator family. In GST pull-down assays, researchers identified regions of the AF-1 that are important for binding to GRIP1 and SRC1α.^[10]^ Through NMR and circular dichroism (CD) studies, it has been shown that the NTD of both ERα and ERβ is unstructured in solution. However, the TATA-box binding protein (TBP) has been demonstrated through surface plasmon resonance and CD to bind to the NTD of ER*α*, suggesting that upon TBP binding, the NTD undergoes structural changes, as observed by a reduction in random coil content.^[11]^ Other nuclear receptors have shown that their NTDs participate in coregulator interactions. The NTD of the AR has been shown to interact with SRC1 in a ligand-dependent manner.^[12]^ It has also been observed that the NTD of rat MR is capable of binding to p300.^[13]^ The NTD of the nuclear receptor Nurr1 has also been shown to interact with SRC1 through the AF-1 region.^[14]^

In addition to interactions with coregulatory proteins, studies have also provided evidence of interdomain interactions in nuclear receptors. For instance, in the AR, the AF-1 domain was found to contain coregulator-like interaction motifs, such as FXXLL and WXXLF, which participate in androgen-dependent NTD and AF-2 binding (for FXXLL). The WXXLF sequence binds in regions outside the AF-2 in the LBD.^[15]^ In the MR, the NTD has been reported to participate in interdomain interaction with the LBD as a result of strong agonist binding.^[16]^ In the glucocorticoid receptor (GR), the N-terminal region of the coactivator TIF2 binds to the NTD/AF-1 region of the receptor, leading to conformational changes in the receptor.^[17]^ In the progesterone receptor (PR), interdomain contacts between the AF-1 and AF-2 regions were observed when the receptor was not bound to DNA.^[18]^ These studies have presented evidence that the NTD of nuclear receptors serves a function in the intrinsic activity of their receptors, whether this is through interacting with coregulator proteins or participating in interdomain NTD-LBD contacts.

We previously explored interdomain contacts in FXR^[19]^ and found that interactions between the DBD and LBD are ligand-dependent and hinge-dependent. The role of the NTD in FXR has not been fully explored, specifically whether it participates in interdomain contacts and/or interacts with coregulators. Using luciferase reporter assays, mammalian two-hybrid assays, and molecular dynamics simulations, we provide evidence for the role of the NTD in FXR by elucidating its interdomain contacts and interactions with coregulators. We show that an SENLF motif in the NTD mediates the observed interactions. As this is the first report of an SXXLF NR-interacting motif, these findings suggest that there may be undiscovered motifs involved in critical NR functions.

## Results

### The FXR NTD interacts with other FXR domains

The NTD of nuclear receptors is known to possess ligand-independent activation function. In FXR, it is not known how the absence of the NTD impacts transcriptional activity. To assess the impact of the NTD on FXR function, we generated a construct with the NTD removed (ΔNTD) (**Fig. 1A)**. Using two luciferase reporter constructs driven by FXREs: IR1-*luc* and PLTP-*luc* (see Methods) we measured and compared transcriptional activity in response to ligand treatment in wild-type FXR (WT-FXR) and ΔNTD constructs. Four FXR agonists were tested: chenodeoxycholic acid (CDCA), fexaramine, obeticholic acid (OCA), and GW4064. Compared to WT-FXR, a significant decrease in response (> 50%) is observed in ΔNTD for all ligands (**Fig. 1B)**. This loss of activity confirms that the NTD is a major contributor to transactivation in FXR. A similar effect was observed in other NRs, such as the human ER^[20]^, AR^[20]^, and PR^[21]^, when the NTD was fully or partially removed.

**Figure 1.**
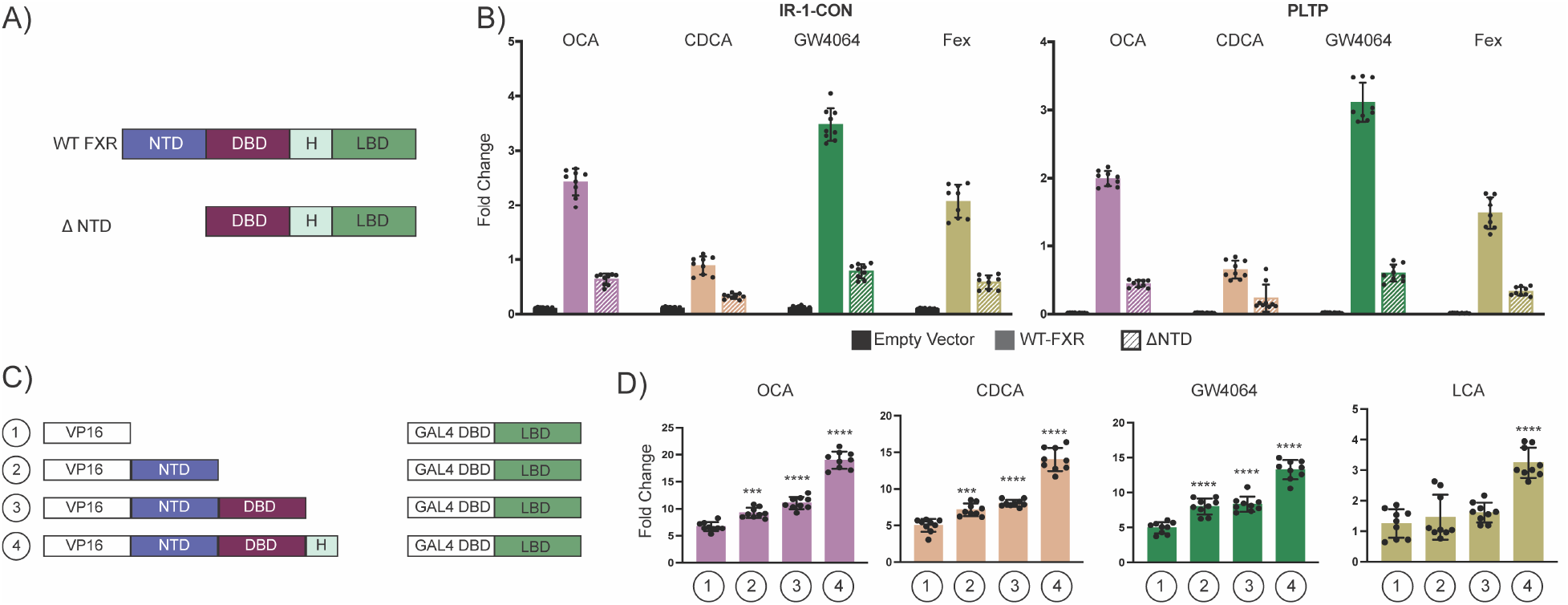
Effect of NTD deletion on FXR activity. A, Schematic representation of WT-FXR and ΔNTD. Domains are colored as follows: NTD in purple, DBD in maroon, hinge (H) in teal, and LBD in green. B, Luciferase assays testing the effect of the NTD deletion on FXR activity. C, Schematic representation of the hybrid constructs used in the mammalian two-hybrid (M2H) assay to assess the interdomain NTD-LBD contacts using the following constructs: NTD, NTD-DBD, and NTD-DBD-H fused to the VP16 individually. The LBD of FXR is fused to the GAL4-DBD, giving GAL4-DBD-FXR-LBD. D, Luciferase assays using the M2H constructs to test interdomain contact between the NTD and LBD. With each additional domain of FXR, the fold change increases relative to the empty VP16 control with GAL4-LBD. Error bars for luciferase assay: standard deviation. B, 2-way ANOVA and D, 1-way ANOVA, p<0.0002 (***), (****) p<0.0001.

**Figure 2.**
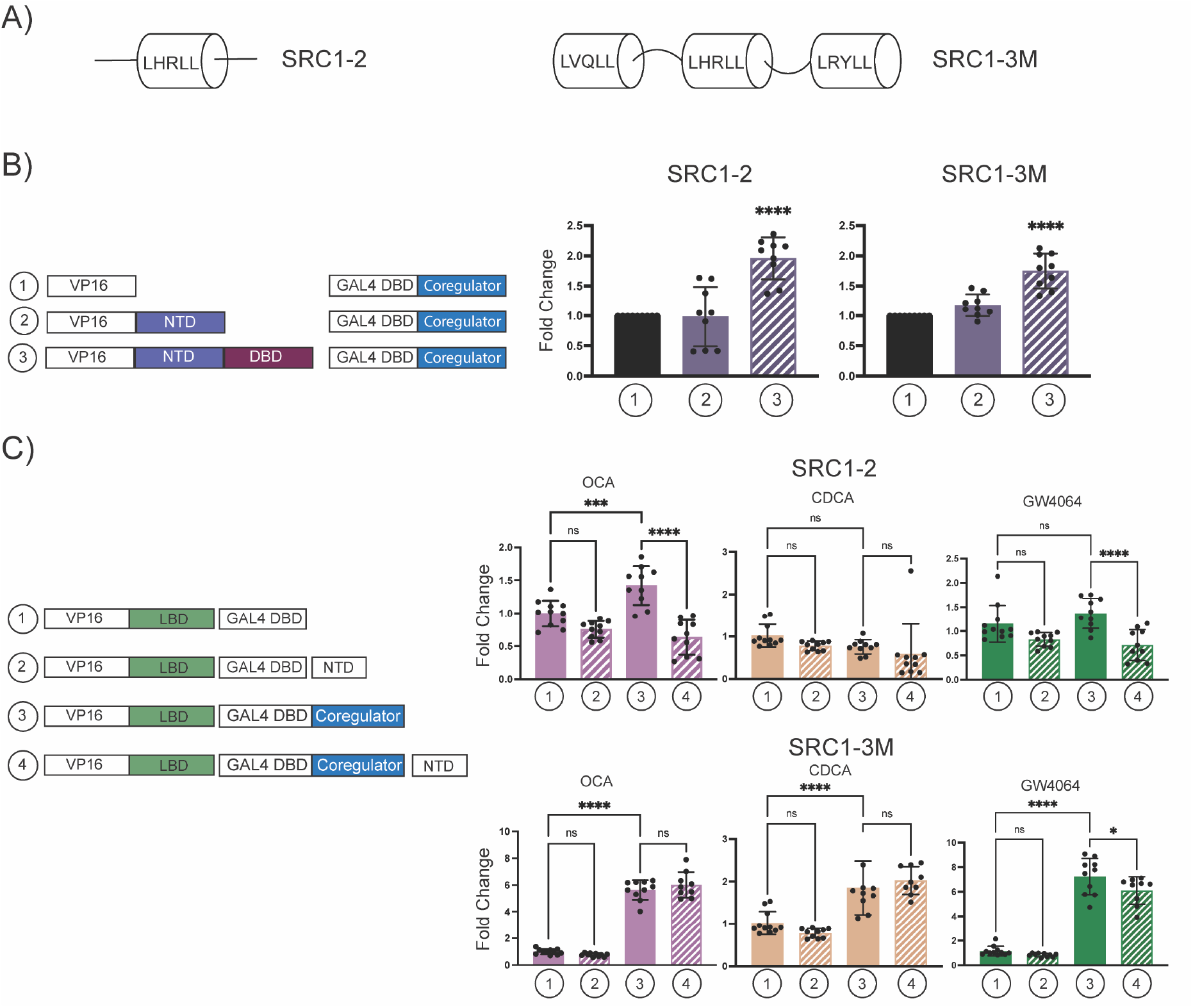
NTD interactions with coregulators. A, Schematic representation of the coregulator fragments used in the M2H system. B, Luciferase assays using the M2H system to assess the interaction between the NTD constructs and SRC1-2 and SRC1-3M. Schematic representations of the constructs used are on the left. A significant increase compared to the control is seen with the NTD-DBD construct for both SRC1-2 and SRC1-3M. C, Luciferase assays using the M2H system to evaluate the effect of the LBD on NTD and coregulator interaction, with schematic representations of the constructs used on the left. A decrease in fold change is observed in the one motif version when the NTD is added, which is not seen in the three-motif version. Error bars for luciferase assay: standard deviation. One-way ANOVA, (****) p<0.0001.

Previous work from our group reported interdomain interactions between the DBD and LBD of FXR. These interactions, observed using an M2H assay, only occurred when the LBD was attached to the hinge domain.^[19]^ We sought to investigate whether a similar M2H assay using the NTD would reveal interdomain interactions with other FXR domains. To determine whether the NTD interacts with the FXR-LBD, two fusion protein constructs were generated: VP16-NTD, with the FXR-NTD fused to the VP16 activation domain, and GAL4DBD-FXR-LBD, with the FXR-LBD fused to the GAL4-DBD **(Fig. 1C)**. When the VP16-NTD and GAL4DBD-FXR-LBD plasmids are co-transfected and treated with FXR ligands, a significant increase in luciferase expression is observed, compared to the control group (empty VP16 and GAL4DBD-FXR-LBD) **(Fig. 1D)**. This observation confirms an interaction between the NTD and LBD. Interestingly, this increase is only observed with FXR agonists CDCA, OCA, and GW4064. The weak agonist LCA did not produce an increase in fold-change over the control **(Fig. 1D)**.

To determine whether other FXR domains influence the NTD-LBD interaction, we created additional constructs: VP16-NTD-DBD and VP16-NTD-DBD-hinge **(Fig. 1C)**. The presence of the DBD led to a small but significant increase in fold-change over the VP16-NTD construct when cells are treated with OCA **(Fig. 1D)**. This suggests that under certain conditions, the DBD may enhance the NTD-LBD interaction. Unexpectedly, the addition of both DBD and hinge to the NTD dramatically increases the interaction with LBD. This is observed for all ligands, including LCA **(Fig. 1D)**. While the cause of the large effect observed with the hinge is not clear, these experiments confirm that the FXR NTD is capable of interactions with the LBD both in isolation and in the intact receptor.

### Peptides from nuclear receptor coregulators interact with the FXR NTD

Previous studies have suggested that the NTD of nuclear receptors can interact with coregulator proteins.^[22–26]^ To explore this possibility in FXR, we again used the M2H system, this time testing for an interaction between VP16-NTD and GAL4DBD-SRC1-2, which contains the SRC1-2 coregulator peptide cloned next to the GAL4DBD. This strategy has been used before to measure nuclear receptor-coregulator interactions.^[27–31]^ Interestingly, we do not observe a significant increase in fold-change over the control when VP16-NTD alone is co-transfected with the coregulator construct. When we test the VP16-NTD-DBD construct, a considerable increase in activity is observed, alluding to an interaction between the NTD-DBD construct and the coregulator fragment. The increased signal is observed for all three ligands tested. To determine whether the NTD can interact with a multivalent coregulator, we created GAL4DBD-SRC1-3M, by cloning the receptor interaction domain of SRC1 into the GAL4DBD plasmid. This fragment, containing 3 LXXLL motifs, interacts with the LBD of nuclear receptors, e.g., PPARγ.^[32]^ In our M2H assay, we observed an increase in fold change between SRC1-3M and the NTD-DBD. Compared with the one-motif version of SRC1, the fold change of SRC1-3M is similar to that of the one-motif version. This could indicate that, among the LXXLL motifs added to the construct, the NTD does not have a greater preference for these additional motifs.

So far, we have confirmed that the FXR-NTD interacts with both the LBD and coregulator peptide fragments. As it is well established that coregulators bind the AF-2 region of the LBD, we sought to test putative roles for the NTD in this interaction. We designed an M2H system to assay the LBD-coregulator interaction, using VP16-FXR-LBD and GAL4DBD-SRC1-2 fusion constructs. To determine whether the NTD affects this interaction, we co-transfected an NTD-only construct, generated by cloning the FXR-NTD into pcDNA3.1 (See Methods). We observed that the addition of the NTD results in a decreased fold-change, suggesting that the disordered domain may interfere with LBD-coregulator interaction. This was observed with OCA and GW4064-treated cells, but not CDCA. Interestingly, when we used the SRC1-3M coregulator motif (GAL4DBD-SRC1-3M), no effect on the LBD-coregulator interaction is observed. This result may indicate that multivalent binding of coregulator motifs is facilitated by SRC1-3M, allowing simultaneous binding to both NTD and LBD.

### A novel SXXLF motif in the FXR NTD mediates interactions with the LBD

Due to the disordered nature of nuclear receptor NTDs, no experimental structural studies have resolved the domain, precluding the ability to generate a structural understanding of how it interacts with other domains. Similar to what was observed in the androgen receptor, we hypothesized that motifs in the FXR-NTD could mediate interdomain and coregulator interactions. We analyzed the NTD sequence in search of known motifs: LXXLL, WXXLF or FXXLF. While none of these were found, the closest motifs identified was SENLF **(Fig. 3A)**. In particular, SENLF is similar to FQNLF in the NTD of the AR, found to participate in agonist-dependent LBD/AF-2 interactions.^[16,20,28,29,33,34]^ To gain insight into the FXR-NTD structure and the predicted secondary structure of these two motifs, we generated models of full-length FXR, including the NTD. Models were generated using AlphaFold3 and, as expected, revealed a highly unstructured NTD, with intact DBD and LBD **(Fig. 3B)**. Interestingly, the SENLF sequence forms a helix and docks near the AF-2 surface of the LBD (**Fig. 3C**).

**Figure 3.**
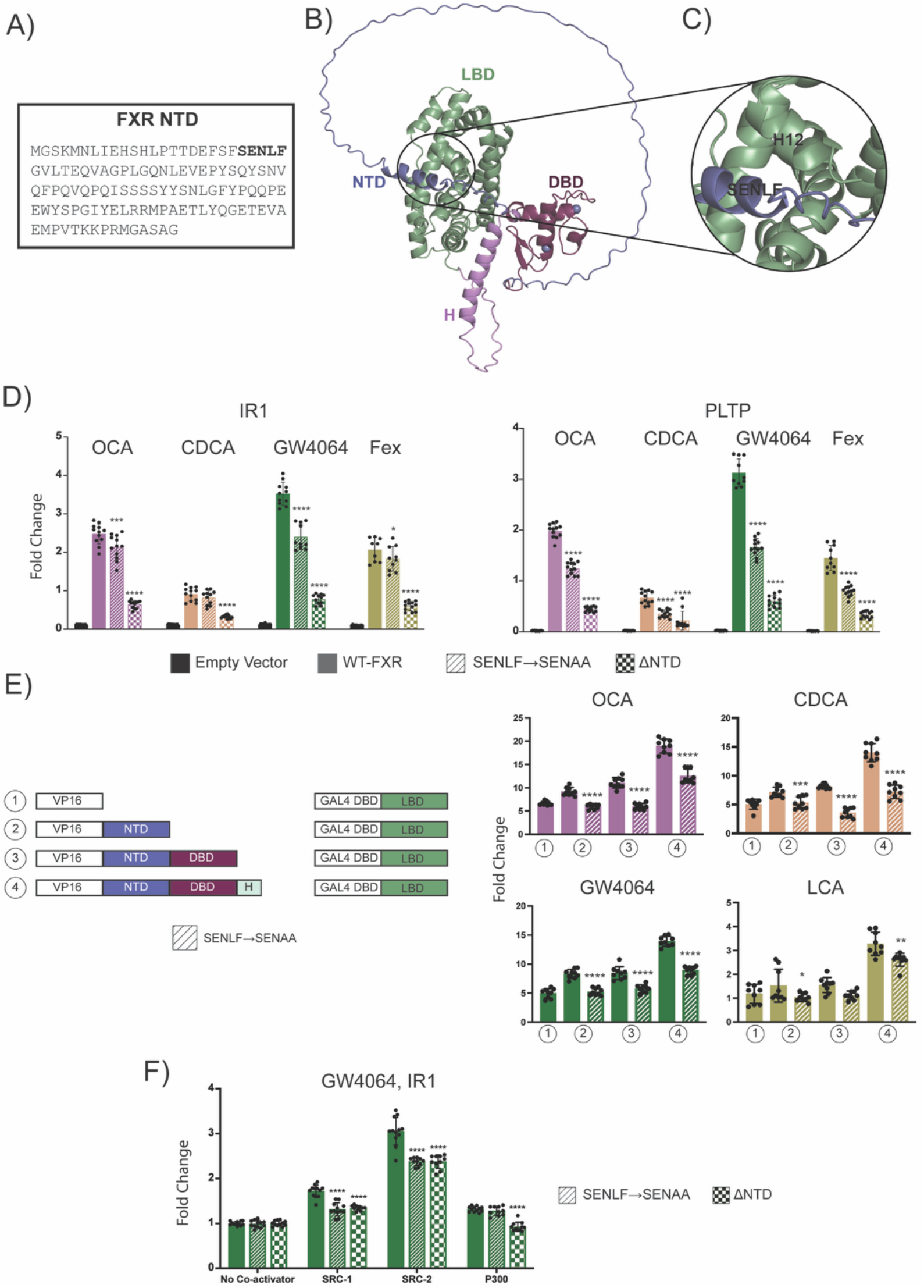
The SENLF motif mediates NTD functions. A, FXR NTD sequence with the SENLF motif bolded. B, AlphaFold3 model of full length FXR. C, The SENLF motif from the NTD is positioned next to H12 in the LBD. D, Luciferase assays evaluating the effect of the SENAA mutation on FXR activity using the IR1-*luc* and PLTP-*luc* constructs. After the mutations, a reduction in fold change was observed. E, M2H assay to assess the effect of the SENAA mutation on the interdomain NTD-LBD contacts using the following constructs: NTD, NTD-DBD, and NTD-DBD-H in the VP16 vector with the LBD in the GAL4-DBD vector. The SENAA mutation is associated with a reduction in interaction for conditions 2-4. F, Luciferase assays evaluating the effect of the SENAA mutation on coregulator utilization normalized to the constructs no coactivator condition. Error bars for luciferase assay: standard deviation. D, 2-way ANOVA and E, 1-way ANOVA, (****) p<0.0001.

To determine whether this sequence is a valid recognition motif in the FXR-NTD, we mutated it to SENAA in our pcDNA3.1-FXR construct and tested the mutant receptor in the luciferase assay. With both reporter constructs (IR1-*luc*, PLTP-*luc*), the SENAA construct significantly decreases fold activation compared to WT-FXR. This is observed in all conditions, except for CDCA with the IR1-luc element, validating the importance of the motif in FXR function. Interestingly, the activity levels of the SENAA mutant are not decreased to the same extent as ΔNTD, indicating that other parts of the NTD may facilitate the receptor’s normal function.

To determine whether SENLF is an LBD recognition motif, we inserted the SENAA mutation into all M2H NTD constructs (NTD, NTD-DBD, and NTD-DBD-hinge) and tested their effect on the LBD-NTD interaction. Compared to WT constructs, the SENAA mutant decreases the fold change, suggesting that the NTD-LBD interaction is disrupted. The decrease was observed in cells treated with OCA, CDCA, and GW4064. In cells treated with LCA, only the NTD-DBD-hinge condition showed a decreased fold change due to the SENAA mutation. These experiments provide strong support for the SENLF sequence as an LBD recognition motif, also validating the positioning of the motif as observed in the AlphaFold predicted structure.

### The SXXLF motif selectively mediates NTD-coregulator interactions

Next, we asked whether the SENLF motif plays a role in coregulator recognition. Using our pcDNA3.1-FXR construct, we quantified the effect of added coregulators on FXR-driven transcription. Our selection of coregulators included SRC-1, SRC-2 (full-length), and p300. When co-transfected with FXR, all three coregulators increased the fold-change, confirming their interaction with FXR **(Fig. 3F)**. We then tested the SENAA mutant and its interaction with coregulators. As a control, we also included the ΔNTD mutant, which should completely abolish NTD-coregulator interactions. As expected, lower fold-change is observed in both SENAA and ΔNTD mutant conditions, compared to the wild-type. Interestingly, for all three variants (WT, SENAA, and ΔNTD), the fold-change increases upon addition of coregulators. This result indicates that all three coregulators retain interactions with FXR regardless of NTD mutations.

To quantify the effect of the mutations on coregulator activity, the data are normalized to the corresponding no-coregulator control for each group **(Fig. 3F)**. The normalization clearly shows that SENAA and ΔNTD mutations have the same magnitude of effect on coregulator activity for SRC-1 and SRC-2. This observation strongly suggests that loss of the SENLF motif recapitulates the effect of ΔNTD on coregulator activity. Conversely, compared to WT-FXR, P300 only shows a decrease in the ΔNTD mutant, suggesting that mechanisms other than SENLF are preferentially used for P300 recognition. Regardless, we note that the NTD mutations do not decrease fold-changes to the level of the no-coregulator controls, supporting the role of the FXR-LBD as the major coregulator interaction surface.

### SXXLF mutation influences interdomain contact and coupling in simulations

To investigate the effect of the SENAA mutation on FXR dynamics, we performed MD simulations of both WT and SENAA FXR models. Six replica 5-μs trajectories were obtained for each monomeric model and combined prior to analysis. A contact analysis was performed to identify differences in contacts between both complexes. Only a handful of minor differences were observed between the two contact maps **(Fig. 4A)**, indicating that no major conformational changes are predicted between WT and mutant monomeric FXR. We hypothesized that conformational effects might be observed in a more physiologically relevant context. Thus, we constructed a model of the DNA-bound FXR-RXR heterodimer, where FXR contains all domains, including NTD, while RXR only contains DBD, hinge, and LBD **(Fig. 4B)**. The RXR-NTD was omitted to remove any confounding effects of a second large, disordered region. The IR1 consensus DNA sequence, which represents the canonical FXR-RXR response element, was used in the model. We generated the SENAA mutant of the heterodimer as well and ran MD simulations (four replicas of length 1.5 to 3.5 μs for each complex). Contact maps reveal clear differences in the FXR-NTD region of both complexes **(Fig. 4C)**. Extensive contacts are observed between amino acids 1-30 of the NTD (containing SENLF) and the C-terminal region of the LBD (containing H12) in the WT complex. These contacts are absent in the SENAA mutant.

**Figure 4.**
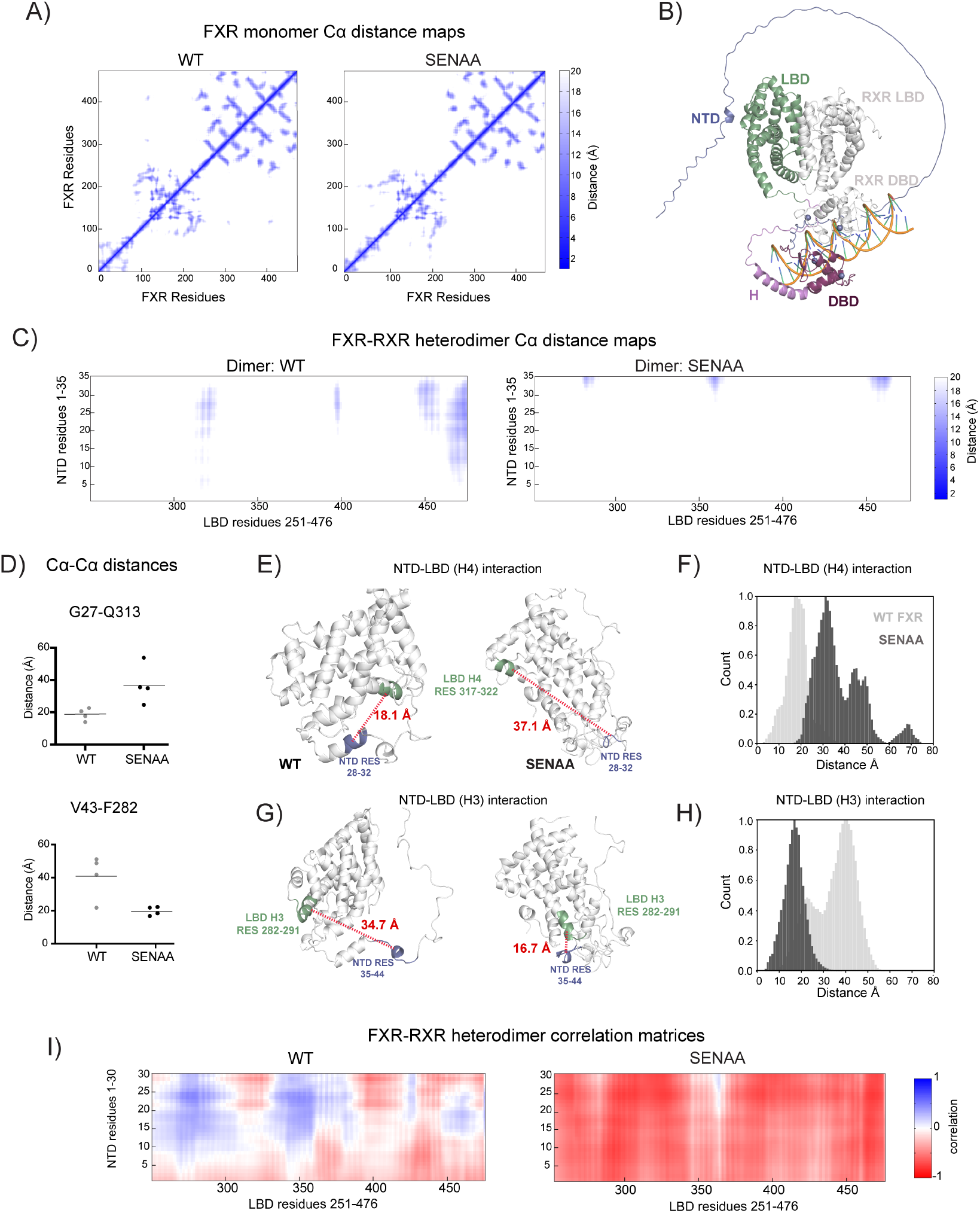
Effect of SENAA mutant on FXR dynamics. A, C-alpha distance maps comparing contacts in monomeric WT and SENAA FXR mutant. B, Alphafold3 model of the FXR-RXR heterodimer, with FXR NTD included. C, C-alpha distance maps comparing contacts in WT and SENAA FXR-RXR heterodimers. D, C-alpha distance between residue pairs G27-Q313 and V43-F282, illustrating decrease and increase in inter-residue distance upon SENAA mutation. E, NTD-LBD (H4) interaction which weakens in the SENAA mutant. F, Histogram summarizing the NTD-LBD(H4) C-alpha distances which are weakened in the SENAA mutant. G, NTD-LBD (H3) interaction which is enhanced in the SENAA mutant. H, Histogram summarizing the NTD-LBD(H3) C-alpha distances which are enhanced in the SENAA mutant. I, Correlation matrices comparing differences in NTD-LBD correlation between WT and SENAA FXR-RXR heterodimers.

To identify the most significant conformational differences between isoforms, we measured average Cα-Cα distances between all FXR residues, and comparing by t-test, determined which inter-residue distances were significantly changed (decreased or increased) in the SENAA mutant. To focus on differences that constitute a major conformational change, we only analyzed residue pairs satisfying two criteria: 1) average inter-residue distance is < 20 Å in either the WT or mutant complex, and 2) the change in inter-residue distance is > 15 Å. Two examples are provided between G27-Q313 and V43-F282 **(Fig. 4D)**. The average G27-Q313 distance changes from 18.7 Å in FXR to 37.2 Å in the SENAA mutant, while the V43-F282 distance decreases from 40.8 Å in FXR to 19.8 Å in the mutant. By our criteria, both displacements represent significant conformational movements as a result of the mutation.

We highlight the region of the largest increase and decrease in inter-residue displacement. The largest increases in inter-residue distance upon mutation is observed between the NTD and LBD H4 **(Fig. 4E, left)**. In the WT complex, we observe that the NTD is positioned proximally to H4, with an average distance < 20 Å. As H4 is part of AF-2, this positioning reflects a close interaction between the NTD and AF-2 regions. Specific NTD residues involved in this conformation are residues 28-33, which lie adjacent to the SENLF motif (22-26). In contrast, in the SENAA mutant, residues 28-33 are more distant from the H4, with average distance > 35 Å **(Fig. 4E, right)**. This change in distances is summarized in a histogram (**Fig. 4F)**. The largest decreases were observed between NTD residues 35-48 and H3 of the LBD (**Fig. 4G)**. The average distance between these regions shifts from ∼ 34 Å in WT FXR to under 20 Å in the SENAA mutant (summarized in **Fig. 4H**). These large shifts in the position of the NTD highlight a significant conformational rearrangement that occurs in response to the mutation.

To characterize how the SENAA mutation potentially influences allosteric coupling between the NTD and LBD, we measured correlation between amino acids 1-30 of the NTD which contain the SENLF motif in the NTD and the LBD **(Fig. 4I)**. A striking difference is observed in correlation patterns. While residues 1-30 of the SENAA mutant complex are anticorrelated with the entire LBD, several regions of positive correlation are observed in the WT complex. This includes the region beginning from H10 up to H12. This result suggests that the two amino acid mutations in the NTD dramatically alter conformation as well as interdomain coupling.

## Discussion

To define the roles of the FXR-NTD, we first asked whether it interacts with other domains. Our data confirms these interdomain interactions. This result is not unexpected, based on precedents in the literature. We observe that adding the DBD and the hinge enhances interdomain interactions. While the reason for this enhancement is not obvious from our experiments, previous reports on the PR indicate that the NTD undergoes structural stabilization when attached to the DBD, also playing roles in DNA binding.^[35]^ The presence of additional FXR domains may impart increased structure to the disordered NTD, enhancing its interactions with other domains or binding partners. Biophysical experiments such as CD or NMR will be used in future experiments to test the hypothesis that additional domains increase the secondary structure of the NTD.

We show that the FXR-NTD can interact with coregulators, as determined by our M2H assay. When a coregulator construct containing one SRC1 LXXLL motif was transfected with the LBD and NTD, we observed a reduction in the interaction between LBD and the coregulator, suggesting a competition between the NTD and LBD for coregulator interaction. Use of the SRC1-3M construct allowed interaction with LBD to be observed, even with co-transfection of the NTD. This result strongly supports multivalent interactions between the three LXXLL motifs on SRC1-3M, FXR-LBD, and NTD. As most coregulators have multiple LXXLL motifs, it is likely that they can simultaneously interact with LBD and NTD.^[36–38]^

We have validated the SENLF motif as a coregulator interaction motif. SXXLF represents a novel recognition motif, with WXXLF and FXXLF from AR being the first time that non-canonical LXXLL coregulator motifs were identified in a nuclear receptor.^[39,40]^ We have now expanded this and have established that SENLF is another recognition sequence for FXR. Non-canonical LXXLL motifs were identified in the orphan nuclear receptor DAX-1, which are involved in repressing the activity of other receptors SF-1 and LRH1.^[41]^ Unveiling this new motif in FXR allows us to probe more deeply regarding how FXR selectively recruits certain coregulators and/or binding partners. For instance, do different motifs preferentially interact with specific coregulators? Our findings also suggest that the full set of NR recognition motifs is not yet known. The identification of SXXLF and related motifs in nuclear receptors and other proteins may drive the discovery of novel NR-interacting partners.

But interestingly, SENLF alone does not capture the full functional contribution of the NTD. We observed this because the SENAA mutant did not reduce FXR transcriptional activity to the same extent as the ΔNTD mutant **(Fig. 3D)**. This finding strongly implicates other regions of the NTD in coregulator interactions and/or crucial NR functions. Future work to identify additional motifs in the NTDs (e.g. by segment deletions, alanine scanning, etc.) as well as elucidate the domain’s mechanism of action will be required.^[42]^

In summary, our work confirms two roles for the FXR-NTD: interactions with the LBD and recognition of coregulators. This work highlights the complexity of NR-coregulator interactions, as we observed that the ΔNTD but not the SENAA mutation impaired recruitment of the P300 coregulator. This finding strongly hints at coregulator specificity of the motif, suggesting that SENLF may preferentially interact with SRC-1 and SRC-2 but not P300. The underlying cause for this phenomenon will be explored in future studies. While the LBD appears to be the most essential factor for binding coregulators, it remains possible that some coregulators prefer the NTD. Our future studies will test a larger swarth of coregulators to understand their partitioning between the two domains.

## Methods

### List of primers

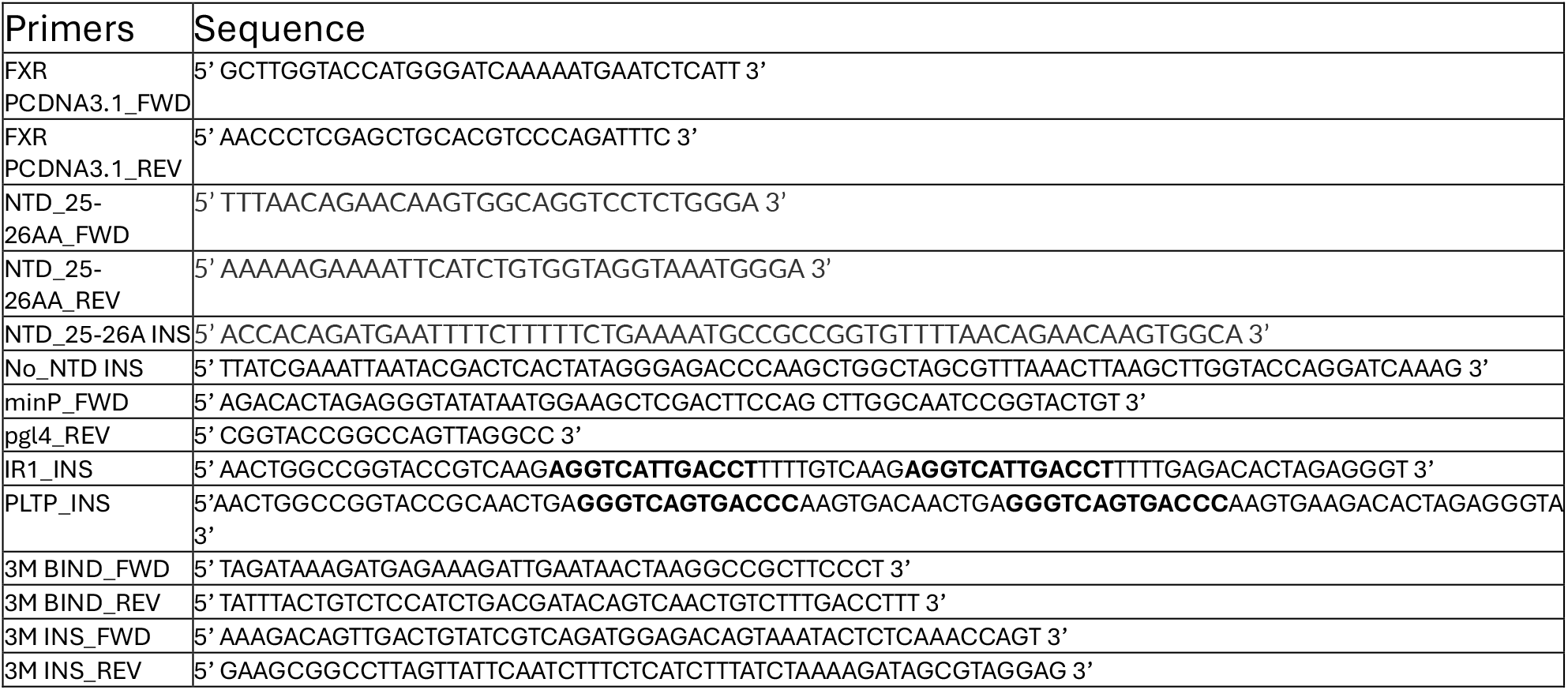

### Cloning/constructs

Human FXRα (NR1H4, UniProt Q96RI1-1) was cloned into the pcDNA3.1(+) expression vector using forward and reverse primers that contained the restriction sites for EcoRI and HindIII. The ΔNTD was made by using PCR to amplify the region of FXR after the NTD and then using this insert (ΔNTD-INS) to assemble the plasmid with a start codon added to the DBD. The constructs for the luciferase reporter assays were made by modifying pGL4.31(luc2P/GAL4UAS/Hygro) with two copies of the FXRE and the minimal promoter upstream of the firefly luciferase gene. pGL4.31 was first digested by Dpn1-HF and HindIII-HF and PCR amplified with primers (minP_FWD and pgl4_REV) to append the minimal promoter. Then, two copies of the FXRE sequence IR1 and PLTP were inserted using Gibson assembly.

GenScript mainly synthesized the constructs used in the mammalian two-hybrid (M2H) protein-protein interaction assay. The LBD of FXR (residues 245-476) was inserted after the GAL4-DBD in the pBIND vector (Promega) to give GAL4-FXR-LBD, and it was also inserted downstream of the VP16 gene in the pACT vector (Promega) to give VP16-LBD. For the NTD (residues 1-119), NTD-DBD (residues 1-196), and NTD-DBD-H (residues 1-244) constructs, they were all individually cloned into the pACT vector downstream of the VP16 gene to give rise to VP16-NTD, VP16-NTD-DBD, and VP16-NTD-DBD-H. For the coregulators used in the hybrid assay, the nuclear receptor recognition motif of SRC1 (residues: LTERHKILHRLLQEGSPSD) was inserted at the N-terminus of the GAL4DBD, yielding GAL4-SRC1-2. The GAL4-SRC1-3M construct was generated by cloning residues 620-759 (Human NCOA1, SRC1α, Uniprot Q15788-1) into the pBIND vector downstream of the GAL4 gene using Gibson assembly.

### Luciferase assays

#### Cell lines and cell culture

Cell reporter assays were performed with HeLa cells (American Type Culture Collection) grown in Dulbecco’s Modified Eagle Medium (Thermo Fisher Scientific) with 10% fetal bovine serum and 1% L-glutamine and incubated at 37 °C in a humidified atmosphere of 5% CO_2_.

#### Transfection and dual-luciferase assay

Luciferase assays and transfections were performed as we have previously described.^[43]^ HeLa cells were seeded at 10,000 cells/well in 96-well, clear flat-bottom cell culture plates. Cotransfection was performed with 5 ng of the expression plasmid pcDNA3.1-FXR, 50 ng of the pGL4.31-derived plasmids containing the luciferase gene downstream of the FXRE, and 1 ng of renilla luciferase (pRL-CMV). The FXREs used were IR1 (inverted repeat spaced by one nucleotide) sequences, the IR1 consensus sequence (AGGTCAtTGACCT), and the PLTP IR1 sequence. The FXRE plasmids used in this assay contained two copies of their respective response element sequences. Transfection was repeated at least in triplicate with Fugene HD (Promega). In assays with added whole coregulator, 10 ng of plasmids containing pSG5-SRC-1, pcDNA3.1-SRC-2, and pDEST26-p300. (Coregulator plasmids from Dr. Eric Ortlund (Emory University)).

Twenty-four hours after transfection, cells were treated with DMSO (final concentration of 0.37%) or with test ligands to a final concentration of 10 µM or 100 µM. After 24h incubation, Firefly and Renilla luciferase activity was assayed using the Dual Glo kit (Promega) on a SpectraMax iD5 plate reader. Data is represented as a fold change, which is the Firefly signal over the Renilla signal, normalized to the DMSO-treated empty vector controls unless otherwise specified in the figure caption.

Data from the luciferase assays are presented here as the mean±standard deviation (SD) of three biological replicates. *P*-values were calculated using two-way ANOVA with Dunnett’s multiple comparisons test to evaluate statistical significance between mean comparisons. One-way ANOVA was also used for data when comparing means across multiple groups while focusing on a single factor. Significance levels are as follows: p> 0.1234 (ns), p <0.0332 (*), p<0.0021 (**), p<0.0002 (***), p<0.0001 (****). Analysis was performed using GraphPad Prism (v. 10.6.0).

#### Mammalian two-hybrid assay

For this assay, the instructions from the CheckMate M2H System (Promega) were used. Transfection was performed using equal amounts (5ng) of each pBIND and pACT construct, along with 50 ng of the reporter plasmid pG5luc, which contains the firefly luciferase gene downstream of the activating response element for the GAL4 DBD. Controls were included in wells with empty pACT or pBIND vectors. Treatment of cells, measurement of luciferase activity, and data processing were performed as described in the previous section.

### Molecular dynamics simulations

FXR models for monomeric (NTD+DBD+H+LBD+DNA) and heterodimer (FXR NTD+DBD+H+LBD and RXR DBD+H+LBD+DNA) forms of the receptor were generated using AlphaFold 3.^[44]^ For the monomeric versions of WT and SENAA FXR, six 5-µs replicate simulations were carried out. The protein models were parameterized using the tLEaP program using the ff14SB protein force field.^[45]^ The complexes were then solvated in an octahedral TIP3P water box with a 10 Å buffer and a sodium chloride salt concentration of 150 mM to mimic physiological conditions.^[45]^ Simulations were minimized, heated, equilibrated, and run with Amber18/20/22.^[46,47,52]^ Simulation equilibration and production were achieved with GPU acceleration.^[48]^ Four minimization steps were completed, each consisting of 5000 steepest descent steps followed by 5000 conjugate gradient steps. The minimization steps are as follows: 1) 500 kcal/mol Å^2^ restraints on solute atoms, 2) 100 kcal/mol Å^2^ restraints on solute atoms, 3) 100 kcal/mol Å^2^ restraints on the ligand, Zn^2+^ ions, and DNA present, and 4) no restraints on the system. In these instances, solute does not refer to sodium chloride ions in solution. After minimization, the system was heated with a 100 ps *NVT* simulation with all atoms under 5 kcal/mol Å^2^ of restraints. Then, pre-equilibration was completed in three 10 ns steps: 1) 10 kcal/mol Å^2^ restraints on solute atoms, 2) 1 kcal/mol Å^2^ restraints on solute atoms, and 3) 1 kcal/mol Å^2^ restraints on the ligand, Zn^2+^ atoms, and DNA present. All simulations used SHAKE with a 2 fs time step.

The simulations for the heterodimer WT and SENAA-FXR complexes were run on the Anton 3^[49]^ supercomputer. The complexes were solvated in a cubic box with a NaCl buffer of varying size, ensuring a total number of atoms exceeding 100,000. Heating, minimization, and pre-equilibration were performed as described above. Four replicas of lengths 1.5 to 3.5 µs were generated for each complex and combined prior to analysis. The heating, minimization, and equilibration for classical MD simulation follow the protocol previously published by our group. Cα-distance maps and correlation matrices were generated using CPPTRAJ.^[50,51]^

## Acknowledgments

This work was supported by an NSF CAREER award CAREER: 2144679 (CDO), a National Institutes of Health award DP2-GM149753 (to CDO), and an NIH T32GM152354 grant (PV)

